# Progressive Actomyosin Remodeling Defining a Structured Podocyte Injury Response in Nephrotic Syndrome

**DOI:** 10.64898/2026.10.07.757076

**Authors:** Ewa Langner, Karla Parra, Shumeng Jiang, Sanjay Jain, Moe R. Mahjoub, Guy M. Genin, Jeffrey H. Miner, Hani Y. Suleiman

## Abstract

Nephrotic syndrome, the leading cause of chronic kidney disease worldwide, is associated with injury and eventual loss of podocytes, the specialized epithelial cells that form the final layer of the kidney’s filtration apparatus. The progression from injury to loss is largely irreversible. However, some forms of nephrotic syndrome undergo remission, suggesting that injured podocytes may retain a capacity for repair, the structural basis of which remains unknown. Podocytes respond to injury by the stereotypical morphological change known as foot process effacement (FPE), which can be visualized using transmission electron microscopy. Using modified ultrastructure expansion microscopy (U-ExM) protocol combined with super-resolution microscopy, we discovered that injured podocytes exhibit a three-stage progression of FPE maturation, with kinetics determined by the degree of actomyosin contractility. Through longitudinal study of adriamycin-induced nephropathy in mice, we identified three sequential stages of actomyosin reorganization within the sarcomere-like structures (SLSs) that define the injury response: focal infiltration of myosin IIA into synaptopodin-rich foot processes (Stage 1), disordered intermingling of synaptopodin/myosin IIA (Stage 2), and the formation of highly organized periodic contractile arrays oriented perpendicular to the direction of capillary blood flow (Stage 3). Longitudinal quantification revealed predictable temporal dynamics, with mature Stage 3 predominating by day 7-9 post-injury. To determine whether this sequence is conserved in human disease, we analyzed kidney biopsies from patients with minimal change disease, focal segmental glomerulosclerosis, membranous nephropathy, and diabetic nephropathy. The same three stages were present across all four diseases, and their distribution tracked disease course: acute podocytopathies exhibited heterogeneous early-stage patterns, whereas chronic injuries converged on uniform, mature SLS organization. These findings demonstrate that podocyte injury is a graded, potentially reversible process rather than a binary switch. They identify structured actomyosin remodeling as a conserved wound response across mechanistically distinct kidney diseases and suggest that therapeutic strategies targeting specific stages of this progression could alter the trajectory of diverse glomerular diseases.

## Introduction

Podocyte injury is ubiquitous across all nephrotic syndromes, the kidney diseases that collectively affect millions of people and remain incurable once kidney filtration is compromised (Levey and Coresh 2012, Rosenberg and Kopp 2017). Podocytes are terminally differentiated epithelial cells whose interdigitating foot processes and slit diaphragms are crucial to maintain the glomerular filtration apparatus (Miner 2012, Butt, Unnersjö-Jess et al. 2020). Because mature podocytes have limited capacity for self-renewal, injury that leads to foot process effacement (FPE) and detachment from the glomerular basement membrane (GBM) is largely irreversible, regardless of whether the cause is autoimmune, metabolic, genetic, or hemodynamic (D’Agati 2008, Schell and Huber 2017). This irreversibility has led to a prevailing view of podocyte injury as a one-way switch: once cytoskeletal architecture is disrupted, the cell is committed to dysfunction or loss.

However, not all injured podocytes die or detach. Clinical observations have long noted that some nephrotic syndromes, notably minimal change disease (MCD), can reverse, implying that podocytes retain a latent repair capacity (Rosenberg and Kopp 2017). A key unknown has been the mechanistic foundation of this healing capacity, and evidence that injured podocytes mount an organized cytoskeletal response rather than simply detaching. Recent super-resolution microscopy studies of injured podocytes identified sarcomere-like structures (SLSs), periodic assemblies of non-muscle myosin IIA, synaptopodin, and α-actinin 4 that resemble the contractile units of striated muscle (Suleiman, Roth et al. 2017). These can be seen in an *ex vivo* culture system with micropatterned substrates that reconstitute the podocyte microenvironment and are known to be mechanoresponsive to substrate stiffness and actomyosin inhibition (Jiang, Alisafaei et al. 2022). SLSs also template synaptopodin-positive peripheral protrusions reminiscent of nascent foot processes and promote podocyte adhesion to the underlying substrate (Jiang, Alisafaei et al. 2022). Complementary genetic studies show that synaptopodin, although not needed for kidney function in healthy mice, is protective during acute injury, consistent with a functional role for SLS-associated synaptopodin in podocyte resilience (Ning, Suleiman et al. 2020, Ning, Suleiman et al. 2021). These findings establish SLSs as mechanosensitive contractile structures with the hallmarks of an injury response to help maintain podocyte adhesion to the glomerular basement membrane (GBM) but leave open the critical question of whether SLS formation represents a structured, staged program or a stochastic byproduct of cytoskeletal collapse.

This distinction is critical from two perspectives. First, a staged remodeling program would imply that podocyte injury proceeds through defined intermediates. These states might be stabilized, accelerated, or reversed by therapeutic intervention. Precedent for such staged programs exists in striated muscle, where sarcomere assembly proceeds through stress fiber intermediates of increasing periodicity (Fenix, Neininger et al. 2018), and in non-muscle cells, where contractile stress fibers mature through progressive actomyosin reorganization (Chapin, Blankman et al. 2012, Tojkander, Gateva et al. 2012). Second, if the same stages are recapitulated across mechanistically distinct human diseases, SLS morphology could serve as a histological readout of where a podocyte sits in its injury trajectory. This would inform diagnostics, which currently classify generalized “damage” by glomerular scarring pattern rather than by the state of cell repair machinery.

Here, we use ultrastructure expansion microscopy (U-ExM) (Langner, Puapatanakul et al.) combined with N-SPARC and Airyscan super-resolution imaging to study a library of mouse models as well as human tissue to test the hypothesis that remodeling defines a structured podocyte stress response in nephrotic syndrome. Data define the *in vivo* trajectory of SLS formation during adriamycin-induced nephropathy in mice and in biopsies from patients with four major forms of nephrotic disease states: MCD, focal segmental glomerulosclerosis (FSGS), membranous nephropathy (MN), and diabetic nephropathy (DN). We identify three structurally distinct SLS stages that define progressive actomyosin reorganization within podocyte foot processes. These findings define podocyte cytoskeletal disruption as an ordered wound response and provide a structural framework for interpreting podocyte injury across the spectrum of nephrotic kidney disease.

## RESULTS

### SLS Formation Progresses in Parallel with Podocyte Injury in Adriamycin Nephropathy

If SLSs represent a programmed wound response rather than passive cytoskeletal collapse, a prediction is that SLS organization should evolve progressively as injury advances, with structural development increasing in step with the severity of podocyte damage. To test this prediction, we combined U-ExM along with N-SPARC super-resolution imaging on mouse kidney sections collected at multiple time points following adriamycin administration, co-staining for nephrin (to mark intact slit diaphragms), synaptopodin (to identify foot process territories), and non-muscle myosin IIA (to detect actomyosin reorganization) (**Figure 1**).

**Figure 1:**
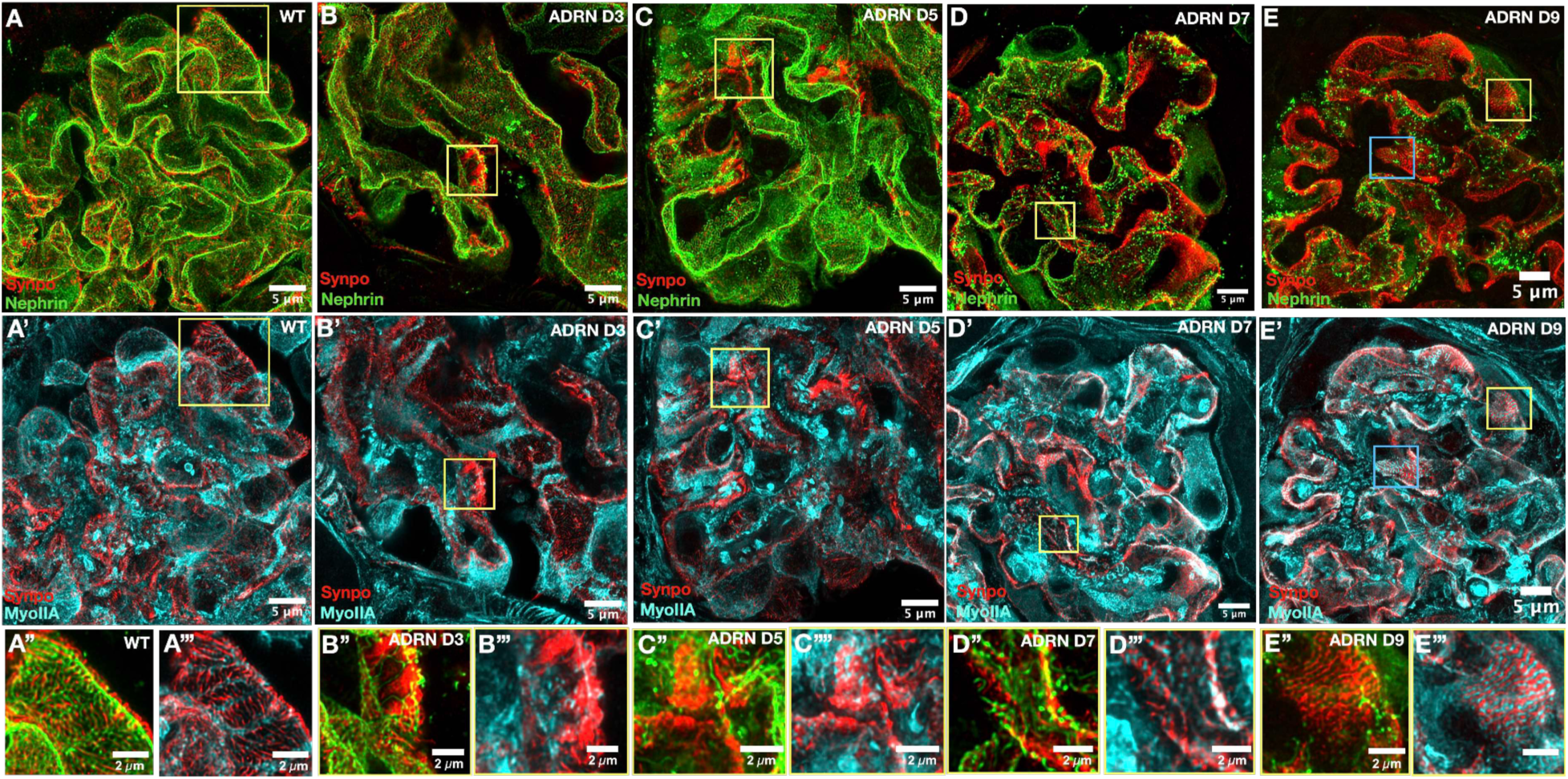
SLS organization increases in parallel with the severity of the podocyte injury in adriamycin nephropathy. U-ExM combined with N-SPARC super-resolution images of paraffin-embedded mouse kidney sections immunostained for nephrin (slit diaphragm marker), synaptopodin (Synpo; foot process marker), and non-muscle myosin IIA (MyoIIA, an SLS marker) at sequential time points following adriamycin injection. At baseline (A, day 0), nephrin staining is continuous along the slit diaphragm and MyoIIA is largely absent from foot processes (A’), indicating intact podocyte architecture with no detectable actomyosin reorganization. At day 3 (B-B’’’), segmental nephrin loss (arrowheads) marks the onset of podocyte injury that persists at day 5 with progressively incorporating MyoIIA patches in those segments (C-C’’’). At day 7 (D-D’’’), nascent SLSs appear as discontinuous, variably oriented MyoIIA/Synpo assemblies within individual foot processes, representing the earliest detectable stage of actomyosin remodeling. By day 9 (E-E’’’), >80% of nephrin signal is lost. SLSs in remaining podocytes have matured into densely packed, well-defined sarcomere-like patterns (E’’ and E’’’). Scale bars, 10 μm.

At baseline (day 0), nephrin staining was continuous along the slit diaphragm and myosin IIA was largely absent from foot processes, indicating intact podocyte architecture with no detectable SLS formation (**Figure 1A**). By day 3, and day 5 after ADR injection, segmental loss of nephrin from the filtration slit marked the onset of podocyte injury (**Figure 1 B-B’’’** and **C-C’’’**, respectively). As predicted, this early injury stage did not present as an overtly disordered cytoskeleton. Instead, myosin IIA appeared within foot processes in discontinuous, variably oriented assemblies recognizable as nascent SLSs but lacking the periodic organization seen in mature contractile structures. This finding is consistent with the hypothesis that SLS formation initiates as a directed cytoskeletal response coincident with, rather than consequent to podocyte shape changes.

By day 7 and 9, podocyte injury had advanced markedly, with >80% loss of nephrin labeling, indicating the reduced density of slit diaphragms due to FPE (**Figure 1 D-D’’’** and **E- E’’’**, respectively). At this stage, SLSs had matured into prominent, well-aligned, and densely packed sarcomere-like bundles. This is in stark contrast to the sparse, disordered assemblies present at day 5. Regions with clustered nephrin were also observed (**Figure 1D** and **E**, Boxes), consistent with complete podocyte detachment and denudation of the GBM. The coexistence of highly organized SLSs in remaining podocytes alongside denuded segments suggests that SLS maturation occurs in cells that are still attached and maintaining contact with the basement membrane, rather than in cells that are already committed to detachment.

Together, these data support the prediction that SLS formation follows a stepwise trajectory that tracks injury severity, progressing from absent, to nascent and disorganized, to highly ordered contractile arrays. This temporal coupling between injury and increasing SLS organization is inconsistent with passive cytoskeletal collapse, which would be expected to produce progressively less organized structures over time. Instead, the results point toward an active remodeling program that becomes more structured as the mechanical challenge to the podocyte intensifies.

### SLS-Enriched Domains Progress from Focal to Near-Global Coverage

The qualitative progression observed in Figure 1 raises a quantitative question: does SLS formation spread gradually across the podocyte’s basal domain as FPE progresses, or does it appear abruptly once a threshold of injury is crossed? These alternatives carry different mechanistic implications. Gradual progression would suggest that individual podocytes initiate SLS assembly locally and then propagate it through their diminishing foot process network as injury intensifies, consistent with a regulated, dose-dependent remodeling process. Abrupt, all-or-none appearance would instead suggest a switch-like response triggered only at a critical level of injury.

To distinguish between these possibilities, we developed a semiautomated image-analysis pipeline to quantify the spatial extent of SLS formation within individual podocytes over time. Total basal podocyte area was delineated by applying Otsu color thresholding to synaptopodin (Synpo) staining on Airyscan maximum-projection images (**Figure 2A**). Within these territories, SLS-containing regions were identified by the co-localization and intertwining of Synpo and non-muscle myosin IIA (MyoIIA) signal, which marks actomyosin bundle formation, and were segmented manually at high spatial precision (**Figure 2B**). The fraction of basal podocyte area occupied by SLSs was then calculated as the percentage of total Synpo-positive area containing SLS-specific Synpo/MyoIIA co-organization.

**Figure 2:**
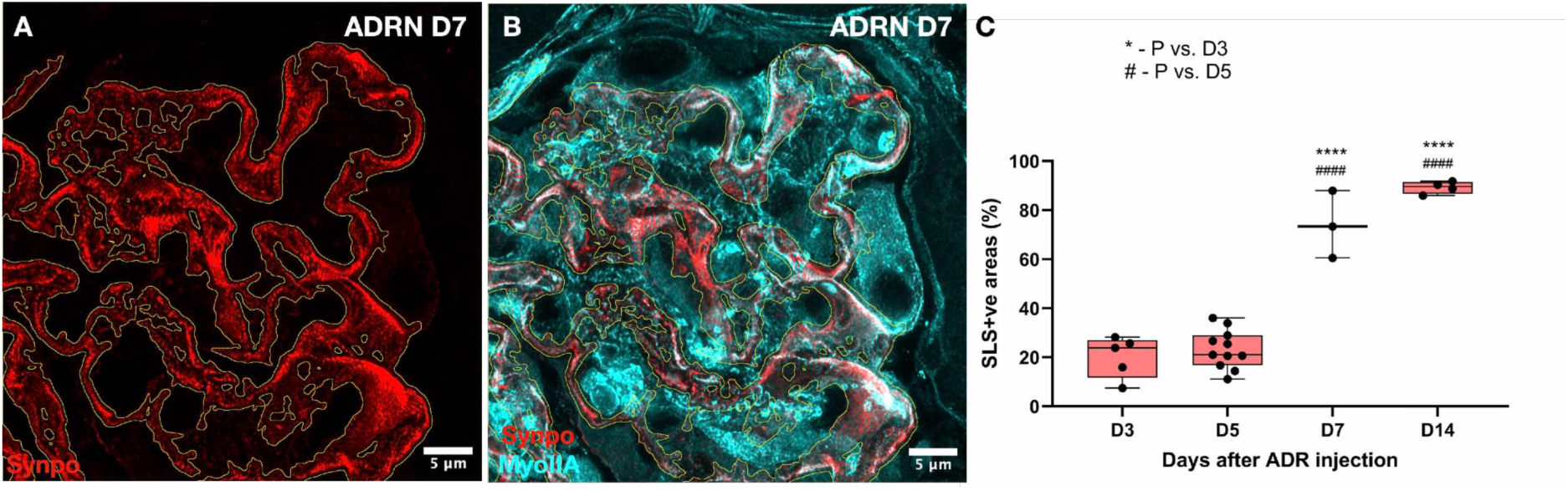
SLS coverage of basal podocyte territory expands progressively from focal initiation to near-complete redistribution. Quantification of SLS spatial extent within podocytes at sequential time points following adriamycin injection. (A) Total basal podocyte area was delineated by applying Otsu color thresholding to synaptopodin (Synpo) staining on Airyscan maximum-projection images. (B) SLS-containing regions, defined by intertwined Synpo and non-muscle myosin IIA (MyoIIA) signal indicative of actomyosin bundle formation, were segmented manually within the Synpo-positive territory. (C) Percentage of Synpo-positive podocyte area occupied by SLSs at each time point. SLSs first appeared as isolated foci at day 3 and expanded progressively, reaching nearly 100% coverage by day 14. The gradual, continuous progression from focal to near-global coverage is consistent with a dose-dependent remodeling process rather than a switch-like structural collapse. Scale bars, 5 μm. **** and #### P<0.0001.

The data strongly favored the gradual progression model. SLSs first emerged as small, isolated foci as early as day 3 following adriamycin injection; this is a time point at which overall podocyte architecture remained largely intact. Over subsequent days, SLS-enriched regions expanded in both density and spatial continuity, spreading from segmental patches to contiguous domains that increasingly spanned the areas of FPE. By day 14, SLSs occupied nearly the entire Synpo-positive domain, representing a transition from limited, localized assembly to near-global coverage of the podocyte actomyosin network (**Figure 2C**).

This progressive spatial expansion has two important implications for subsequent study design. First, it reinforces the conclusion from the qualitative imaging that SLS formation is not a catastrophic structural failure but instead a graded remodeling process whose extent scales with injury severity. Second, the early focal appearance of SLSs at day 3, when nephrin loss is minimal and foot process architecture is still largely preserved, indicates that actomyosin reorganization is among the earliest detectable responses to podocyte injury, preceding the gross morphological changes conventionally used to identify FPE.

### Three Structurally Distinct SLS Stages Define a Sequential Progression of Injury Response

Our discovery of the progressive expansion of SLSs establishes that actomyosin remodeling scales with injury severity, but it does not address whether the internal architecture of SLSs also evolves. If SLS formation reflects a programmed maturation sequence, we would expect to find discrete structural intermediates: instead of a continuum of random configurations, we would expect a small number of reproducible organizational states that appear in a predictable temporal order. Such staging would parallel the well-characterized intermediates of sarcomere assembly in striated muscle, where disordered actomyosin networks condense through progressively more periodic states before reaching mature contractile organization (Fenix, Neininger et al. 2018).

To test this prediction, we performed detailed structural analysis of SLS morphology across all time points in the adriamycin nephropathy model and asked whether individual SLSs could be classified into reproducible organizational categories. Three distinct patterns emerged (**Figure 3A-C**). **Stage 1** was characterized by foot processes in which synaptopodin occupied most of the cross-sectional area, with only sparse, focal infiltration of myosin IIA bundles. In this configuration, which represents the earliest detectable departure from normal foot process architecture, the actomyosin machinery has begun to assemble, but synaptopodin organization remains largely intact and the periodic structure characteristic of mature SLSs has not yet formed (**Figure 3A**).

**Figure 3:**
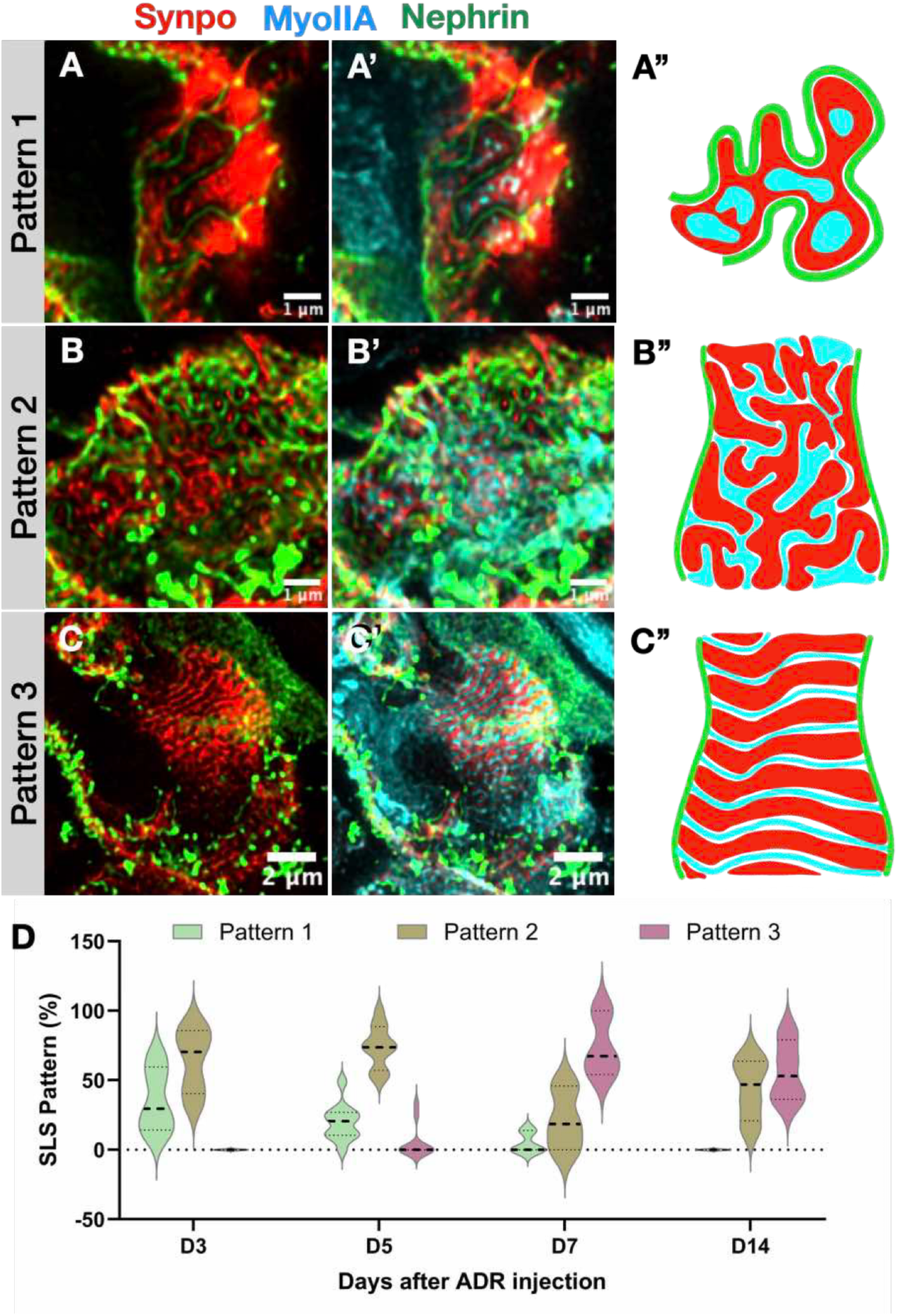
SLS architecture resolves into three discrete structural stages that progress in a stereotyped temporal sequence. Classification and temporal dynamics of SLS organizational states during adriamycin nephropathy. (A) Stage 1: synaptopodin (Synpo) occupies most of the foot process area with only sparse, focal myosin IIA (MyoIIA) infiltration, representing the earliest departure from normal architecture. (B) Stage 2: Synpo and MyoIIA are extensively intermingled in alternating but disorganized domains; residual Synpo-positive bundles persist at the foot process periphery. This transitional configuration reflects active cytoskeletal reorganization that has progressed beyond focal infiltration but has not yet achieved periodic order. (C) Stage 3: highly periodic Synpo/MyoIIA striations are oriented perpendicular to the longitudinal foot process axis, with complete loss of peripheral Synpo bundles. This is the most structurally mature state, resembling the periodicity of sarcomeric contractile units. Schematic drawings (right) depict the organizational features defining each stage. (D) Proportion of each SLS stage at sequential time points following adriamycin injection. Stage 1 dominates at early time points, Stage 2 peaks at day 5, and Stage 3 accounts for over 80% of all SLSs from day 7 onward. The ordered, unidirectional temporal succession is consistent with a deterministic maturation program rather than stochastic structural disintegration. Scale bars, (A-B): 1 μm; (C): 2 µm.

**Stage 2** was defined by a transitional architecture in which synaptopodin and myosin IIA were extensively intermingled in alternating but disorganized condensates. Residual synaptopodin-positive bundles persisted at the foot process periphery, but the clear longitudinal synaptopodin axis of healthy foot processes had been lost. This intermediate state shows neither the focal infiltration of Stage 1 nor the periodicity of Stage 3 and is the structural signature of actively progressing reorganization (**Figure 3B**).

**Stage 3** represented the most advanced configuration: highly periodic, well-aligned Synpo/MyoIIA striations oriented perpendicular to the original longitudinal foot process axis, accompanied by complete loss of peripheral synaptopodin bundles. This is the most sarcomere-like state, with a degree of spatial periodicity that implies coordinated actomyosin self-organization rather than stochastic redistribution (**Figure 3C**).

The distribution of these three stages shifted in a stereotyped temporal sequence following adriamycin injection (**Figure 3D**). Stage 1 predominated at the earliest time points, consistent with the initial, mild perturbation of foot process architecture, and decreased with time progression. By day 5, Stage 2 had become the most prevalent configuration, indicating that the majority of SLS-bearing foot processes had transited through the focal infiltration phase into active reorganization. Beginning at day 7 and continuing through day14, Stage 3 accounted for >80% of all SLSs, reflecting near-complete conversion to the mature contractile phenotype.

This ordered temporal succession across stages is difficult to reconcile with passive cytoskeletal collapse, which would be expected to produce increasingly disordered structures or a mixture of configurations at any given time point. Instead, the data are consistent with a deterministic maturation program in which each stage represents a necessary structural intermediate towards an end point of fully periodic contractile assembly. The existence of such intermediates suggests that SLS maturation could, in principle, be arrested, accelerated, or reversed at specific stages, a possibility we explore in the human biopsy data below.

### SLSs are a Consistent Feature of Human Nephrotic Syndrome Across Mechanistically Distinct Diseases

The adriamycin nephropathy data establish that SLSs mature through a stereotyped three-stage program in a single mouse model of toxin-induced podocyte injury. A strong test of whether this program reflects a fundamental property of injured podocytes, rather than being unique to adriamycin toxicity, is to ask whether the same structures appear in human diseases driven by entirely different pathogenic mechanisms. Nephrotic syndrome provides an ideal test case because its major forms span a wide range of etiologies: autoimmune (MN), metabolic (DN), idiopathic or genetic (primary FSGS), and poorly understood acute podocytopathies (MCD). If SLSs are present across all of these conditions, the implication is that actomyosin remodeling into sarcomeric arrays is a conserved podocyte response to injury itself, independent of the upstream trigger.

We therefore performed super-resolution imaging on kidney biopsy specimens from patients with MCD, primary FSGS, MN, and advanced DN. As controls, we used nephrectomy samples from individuals with negligible proteinuria (**Figure 4A, Mov. 1**). Similar to the method described above (**Figure 2A**), we identified areas that quantify as SLS extents within individual glomeruli, and applied the same analysis pipeline developed for the mouse studies: total foot process area was delineated by standardized color thresholding of synaptopodin staining (**Figure 2A**), and SLS-containing regions were identified by prominent Synpo/myosin IIA intertwining coincident with reduced slit diaphragm density indicated by loss of nephrin (**Figure 4B-H**).

**Figure 4:**
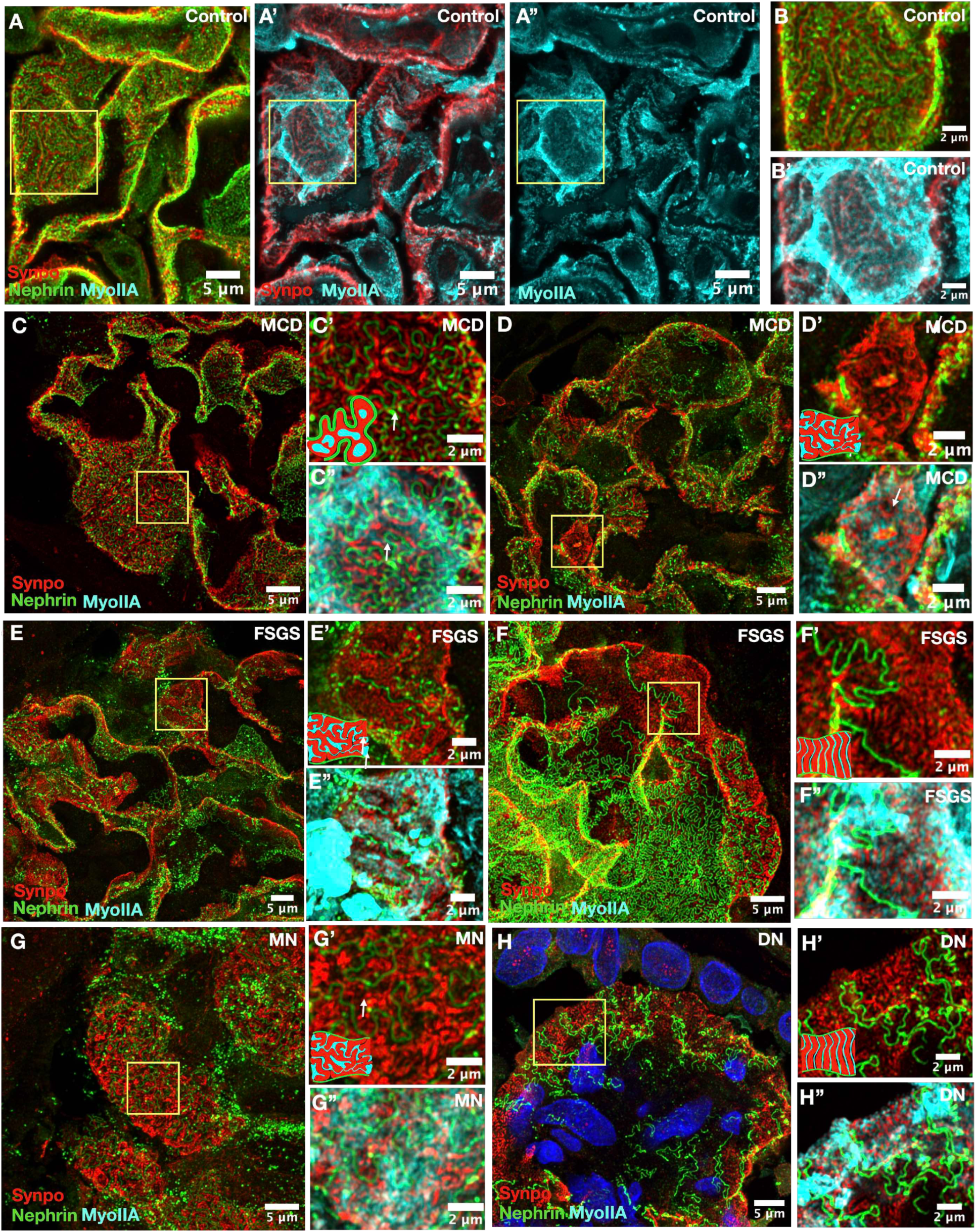
Heterogeneous SLS stages in podocytopathies of various etiologies. Super-resolution en face images of podocyte foot processes of control and nephrotic syndrome biopsies. Control glomerulus (A-A’’ and B-B’) where foot processes display slender, regularly interdigitated morphology with synaptopodin (Synpo, red) forming a central longitudinal axis and minimal non-muscle myosin IIA (MyoIIA, cyan) signal. MyoIIA is confined within the boundaries of podocyte cell body and major processes. Minimal change disease (MCD, C-C’’ and D-D’’) podocytes exabit areas of both stage 1 (C’-C’’) and stage 2 (D-D’’) SLS formation, in different glomeruli of the same patient. Notice the partial preservation of the foot processes, producing a heterogeneous remodeling landscape. Primary focal segmental glomerulosclerosis (FSGS, E-E’’ and F-F’’) exhibiting similar heterogeneity to the one observed in MCD, with progressed stage 2 (E-E’’) and advanced stage 3 (F’-F’’) SLS formations. Membranous nephropathy (MN, G-G’’) foot process broadening and SLS formation are widespread, with prominent stage 2 periodic Synpo/MyoIIA arrangements spanning large contiguous regions. Advanced diabetic nephropathy (DN, H-H’’) uniform foot process expansion with extensive, highly organized stage 3 SLS coverage. Scale bars, 2 and 5 μm.

Maximal projection of super-resolution imaging show en face views of the capillary walls with the interdigitating foot processes and provide a qualitative architectural difference between disease categories (**Figure 4C-H**). In control glomeruli, foot processes displayed a characteristic slender, regularly interdigitated morphology, with synaptopodin forming a central longitudinal axis and minimal myosin IIA signal (**Figure 4A-B**). All nephrotic syndrome samples showed foot process broadening as evident by the nephrin staining (**Figure 4 C, D, E, F, G and H)**, synaptopodin redistribution, and the increased Synpo/MyoIIA intertwining that are the hallmarks of SLS formation. However, the spatial distribution of these changes differed. In MCD (**Figure 4C-C’’, D-D’’, Mov. 2-3**) and primary FSGS (**Figure 4E-E’’, F-F’’, Mov. 4-5**), SLS-bearing foot processes were interspersed among partially preserved regions, producing a heterogeneous landscape of remodeled and intact areas. In MN (**Figure G-G’’, Mov. 6**) and advanced DN (**Figure 4H-H’’, Mov. 7**), remodeling was more uniform, with prominent periodic SLS arrangements spanning large contiguous domains of the foot process area.

SLSs were present in every disease category examined, confirming that actomyosin remodeling into sarcomeric assemblies is not restricted to a single injury mechanism. In control samples, SLSs occupied only 11% of total synaptopodin-positive foot process area (IQR 9-23%), consistent with minimal actomyosin reorganization under physiologic conditions (**Figure 5A**). All four nephrotic syndrome groups showed marked expansion of SLS-enriched regions. In the acute podocytopathies, SLSs accounted for 71% of foot process area in MCD (IQR 44-86%) and 61% in primary FSGS (IQR 43-86%); note that both diseases exhibited substantial inter-patient variability, as some biopsies showed extensive SLS coverage while others retained large regions of SLS-free foot processes. In the chronic diseases, SLS involvement was more pronounced and more uniform: 86% in MN (IQR 83-92%) and 79% in advanced DN (IQR 67-80%), with SLSs distributed diffusely across the podocyte population rather than in the patchy pattern observed in MCD and FSGS (**Figure 5A**).

**Figure 5:**
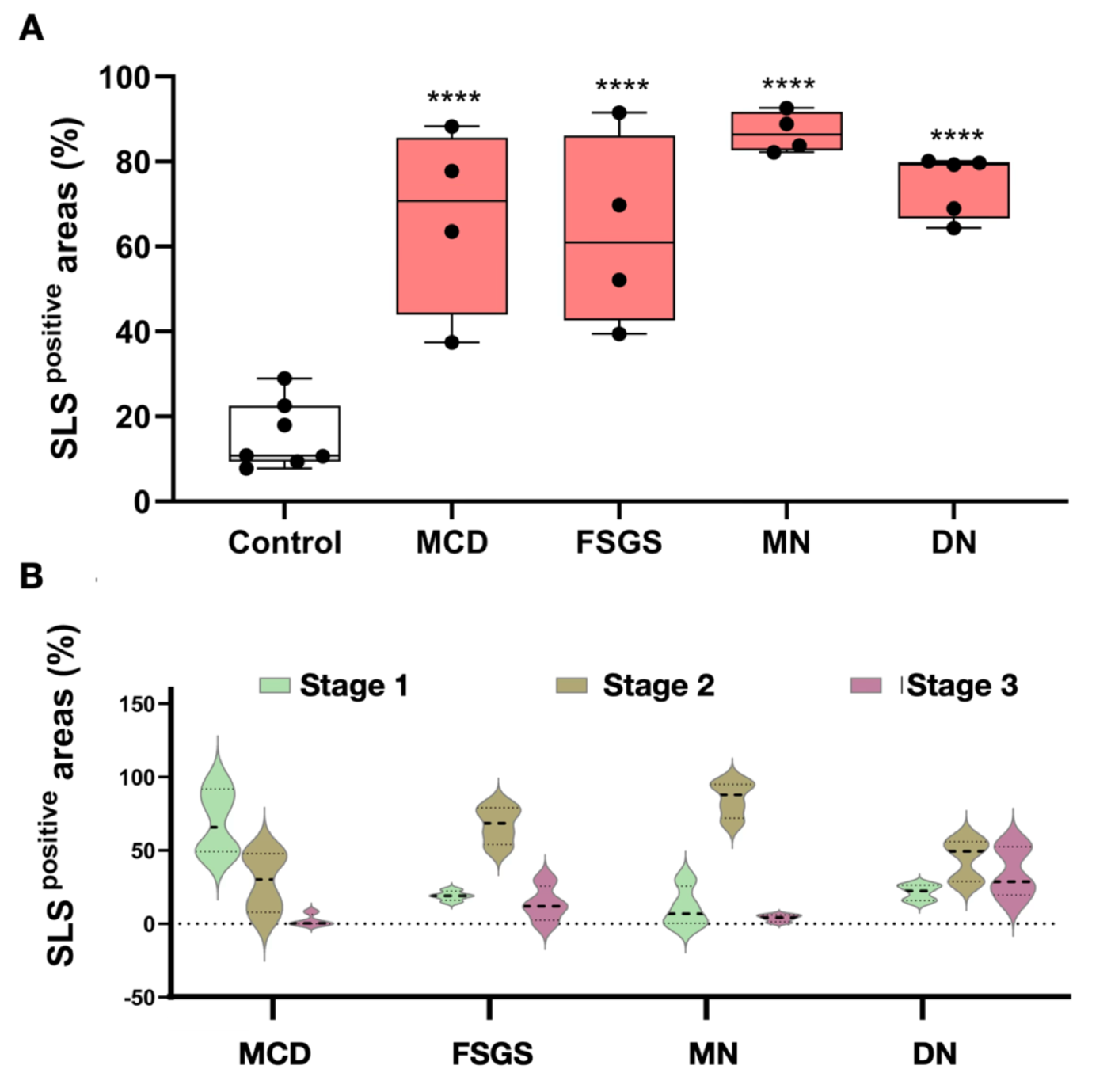
Prevenance of the SLS formation in human glomerulopathies of various nephrotic and their predominant stages. (A) Percentage of Synpo-positive foot process area occupied by SLSs in controls and across four forms of nephrotic syndrome. Controls show minimal SLS involvement (median 11%). Acute podocytopathies (MCD, 71%; FSGS, 61%) show substantial but variable SLS coverage, while chronic diseases (MN, 86%; DN, 79%) show extensive, uniform SLS expansion with less inter-patient variability. The presence of SLSs across all four mechanistically distinct diseases indicates that sarcomeric actomyosin remodeling is a conserved feature of podocyte injury. **** P <0.0001. (B) SLS stage distribution shifts from early to mature configurations across nephrotic syndrome subtypes. Proportion of Stage 1, Stage 2, and Stage 3 SLS configurations in MCD (Stage 1(66%), Stage 2 (30%) and Stage 3 (1%); Primary FSGS Stage 2 (69%), Stage 3 (12%); MN show predominantly Stage 2 (88% and Stage 3 at 4%). Advanced DN exhibits the broadest stage distribution and the highest Stage 3 proportion (29%).

The presence of SLSs across four mechanistically unrelated diseases, spanning autoimmune, metabolic, and idiopathic etiologies, argues that sarcomeric actomyosin remodeling is a general property of the podocyte injury response, not a pathway-specific epiphenomenon. The striking difference in spatial heterogeneity between acute and chronic diseases raises a question that the next section addresses: does this heterogeneity reflect differences in which stages of SLS maturation are present?

### SLS Stage Distribution Evolves Uniformly Across Nephrotic Syndrome Subtypes

The preceding analysis demonstrated that SLSs are universally present across human nephrotic syndromes but differ in spatial extent and uniformity between acute and chronic diseases. We next asked whether these differences reflect distinct distributions of the three SLS maturation stages identified in the mouse model. This question guides translation of the staging framework: if the same three structural states observed in adriamycin nephropathy are present in human biopsies, and if their relative proportions differ in a way that tracks disease course, the implication is that SLS maturation captures where podocytes sit in their injury trajectory, beyond what conventional histopathology provides.

We therefore classified all SLS-containing foot process regions in the human biopsy cohort according to the same three-stage criteria established in mice and quantified the proportion of each stage within each disease category (**Figure 5B**). The distributions all evolved over the same pathway but differed kinetically across diseases. In controls, the minimal SLS formation that was present consisted almost entirely of Stage 1 (93%, IQR 66-100%), with only rare Stage 2 configurations (7%, IQR 0-34%) and no detectable Stage 3. This baseline distribution is consistent with the expectation that healthy podocytes, while capable of low-level actomyosin reorganization, do not progress to more advanced remodeling states in the absence of sustained injury.

MCD, the most acute nephrotic syndrome, was dominated by Stage 1 (66%, IQR 49-92%) with a moderate proportion of Stage 2 (30%, IQR 8-48%) and minimal Stage 3 (1%, IQR 0-6%) (**Figure 5B**). This closely resembles early time points (days 3-5) of the mouse adriamycin model when injury is established but SLS maturation has not yet progressed beyond the initial and transitional phases. The predominance of early-stage SLSs in MCD is notable because it aligns with the clinical behavior of this disease: MCD frequently remits with treatment, suggesting that the podocyte injury state it features may be more readily reversible than that of other nephrotic syndromes.

Primary FSGS showed a marked shift toward later stages, with Stage 1 reduced to 19% (IQR 16-22%), Stage 2 comprising the majority at 69% (IQR 54-79%), and Stage 3 present in a significant minority at 12% (IQR 3-26%) (**Figure 5B**). This was an intermediate distribution in which transitional remodeling predominates, but mature contractile assembly has begun and mirrors the day 5-7 window in the mouse model. This is consistent with the subacute to chronic clinical course of FSGS, in which podocyte injury is more sustained and less readily reversed than in MCD.

MN, a chronic immune-mediated disease, exhibited a distribution strongly enriched for Stage 2 (88%, IQR 72-95%), with markedly depleted Stage 1 (7%, IQR 0-26%) and surprisingly little Stage 3 (4%, IQR 1-6%) (**Figure 5B**). The near-complete conversion from Stage 1 to Stage 2 indicates that virtually all MN podocytes have transitioned into active reorganization. However, the relative scarcity of Stage 3 distinguishes MN, the source of pathology for which subepithelial immune complex deposition versus direct toxic or metabolic stress in DN suggests that SLS maturation kinetics are determined in part by mechanobiological factors.

Advanced DN showed the broadest distribution across all three stages and the highest proportion of mature SLSs: Stage 1 at 22% (IQR 16-26%), Stage 2 at 50% (IQR 29-56%), and Stage 3 at 29% (IQR 20-53%) (**Figure 5B**). The enrichment for Stage 3 is consistent with prolonged, progressive diabetic podocyte injury, for which years of metabolic and hemodynamic stress may underlie SLS maturation to completion. The coexistence of all three stages within individual DN biopsies may reflect the asynchronous onset of injury across the podocyte population, as appears via the focal and segmental nature of diabetic glomerular disease.

Taken together, these data demonstrate that the three-stage SLS classification developed for the mouse model translates directly to human disease, and that stage distribution varies systematically across nephrotic syndrome subtypes. Results support the interpretation that SLS maturation reflects cumulative actomyosin remodeling driven by the duration and nature of podocyte stress. Disease-specific deviations from a simple chronicity gradient, particularly the Stage 2 arrest in MN, indicate that stage distribution encodes information about injury biology beyond what can be inferred from clinical time course alone.

## DISCUSSION

The prevailing understanding of podocyte injury conceptualizes cytoskeletal disruption as a transition from intact architecture to effacement, with severity graded along a continuum but with cytoskeletal consequences considered largely independent of injury trajectory or recovery potential (D’Agati 2008, Schell and Huber 2017). By resolving the internal organization of sarcomere-like structures across time in a mouse injury model and across four mechanistically distinct forms of human nephrotic syndrome, we show that podocyte actomyosin remodeling proceeds through three discrete structural stages in a predictable sequence. At least in adriamycin-induced nephropathy, mature Stage 3 structures dominate by day 7-9 post-injury. This staging pattern is inconsistent with passive collapse and instead supports the existence of an ordered contractile wound response that carries information about both the duration and the nature of injury.

The maturation sequence follows organizational principles analogous to contractile assembly in other cell types: initial motor protein infiltration into a pre-existing scaffold, followed by progressive self-organization into ordered periodic arrays. Cardiomyocyte sarcomeres mature through stress fiber intermediates of increasing periodicity (Fenix, Neininger et al. 2018), and non-muscle stress fibers in fibroblasts undergo analogous progressive ordering of α-actinin and myosin II (Chapin, Blankman et al. 2012, Tojkander, Gateva et al. 2012). A critical difference, however, is that SLS formation/patterns in podocytes could be altered the podocytes’ in response to various mechanical stimuli (Jiang, Alisafaei et al. 2022), implying that staging reflects the ongoing balance between injury-driven remodeling and baseline cytoskeletal organization. This balance could, in principle, be shifted therapeutically.

The conservation of all three stages across MCD, FSGS, MN, and DN argues that SLS maturation is a fundamental property of the podocyte injury response rather than a pathway-specific phenomenon. This is consistent with the idea that podocytes face a common biophysical problem regardless of the upstream trigger: maintaining adhesion to the GBM under filtration forces (Endlich and Endlich 2012). Within this framework, SLSs may function as a generalized contractile adaptation to mechanical stress (Jiang, Alisafaei et al. 2022).

Disease-specific deviations from a simple chronicity gradient also provide mechanistic insight. Despite its chronic clinical course, MN was characterized predominantly by Stage 2 with minimal Stage 3 representation, whereas advanced DN exhibited the highest proportion of Stage 3 structures. This divergence likely reflects differences in the biomechanical context of injury. In DN, progressive GBM stiffening (Wyss, Henderson et al. 2011, Embry, Mohammadi et al. 2016) together with sustained hemodynamic stress may provide the mechanical input necessary to push SLSs through the final maturation stage. In contrast, the complement-mediated and potentially episodic nature of injury of MN (Ronco and Debiec 2015) may lack the sustained directional stimulus needed for consolidation into Stage 3. These findings suggest that SLS stage distribution capture aspects of injury biology not reflected by clinical chronicity alone.

These results also carry diagnostic and therapeutic implications. Current histopathology assessment primarily captures cumulative glomerular damage and does not directly evaluate whether the podocyte repair machinery is active, exhausted, or potentially reversible. SLS staging may therefore provide a complementary readout of cellular remodeling states. The predominance of Stage 1 SLSs in MCD, a disease with substantial remission potential, raises the possibility that early-stage distributions mark injury states with greater reversibility, whereas Stage 3 SLSs in advanced DN may mark podocytes that have reached a mature remodeling state. More broadly, the existence of discrete maturation intermediates suggests a therapeutic axis: interventions that stabilize the transition from Stage 1 to Stage 2 or prevent progression from Stage 2 to Stage 3 could alter the trajectory of podocyte injury. The mechanosensitivity of SLS formation (Jiang, Alisafaei et al. 2022), together with emerging evidence implicating microtubule dynamics in SLS assembly (Suleiman, Roth et al. 2017), highlight potential molecular entry points for such strategies.

For super-resolution imaging, we used both Airyscan and N-SPARK microscopes that provided similar resolution (∼120 nm). However, such resolution was not enough to visualize the detailed organization of SLSs. Therefore, we used the U-ExM, an expansion microscopy protocol previously adopted for both mouse and human kidney tissue that provided a 4-fold isotropic expansion and super-resolution imaging with a conventional diffraction-limited microscope (Langner, Puapatanakul et al.). Combining these two techniques allowed us for comprehensive and efficient analyses of SLSs structures with a nanoscale-resolution with resolution of (∼30 nm). For imaging the SLSs in vivo, we find this approach more practical when compared with other super-resolution modalities such as the stochastic optical reconstruction microscopy (STORM) (Suleiman, Zhang et al. 2013, Suleiman, Roth et al. 2017), the stimulated emission depletion microscopy (STED) (Unnersjö-Jess, Scott et al. 2016) and the structured illumination microscopy (SIM) (Siegerist, Ribback et al. 2017) techniques.

Several limitations should be noted. First, our SLS staging framework is derived from a progressive, and not reversable injury mouse model, and therefore, we cannot establish whether SLSs are reversable or not, and if they are reversable, which stages can fully reverse to the healthy state. Evaluating the progression of SLS assembly/disassembly in a reversable injury, such as nephrotoxic serum injury model (Verma, Venkatareddy et al. 2018), might answer this question. Second, the human data are cross-sectional, and temporal dynamics were inferred through comparisons across diseases with differing expected chronicity rather than from longitudinal observation. Finally, although our classification identifies dominant organizational states, it does not directly assess the contractile function of SLSs at each stage.

## CONCLUSION

Using U-ExM Super-resolution microscopy, we demonstrate that podocyte cytoskeletal remodeling during injury is not passive structural collapse but a conserved, three-stage contractile maturation program. In the adriamycin nephropathy mouse model, these stages progress in a stereotyped temporal sequence, with mature periodic assemblies predominating as injury advances. In human nephrotic syndrome, the same three stages are present across four mechanistically distinct diseases, and their distribution differs systematically in ways that reflect both disease chronicity and injury biology. These findings reframe podocyte injury as a graded, potentially targetable process and establish SLS staging as a new structural framework for characterizing the state of the podocyte wound response. By revealing that injured podocytes retain the capacity for organized cytoskeletal remodeling, and that this remodeling follows a predictable program with discrete intermediates, this work identifies a previously unrecognized window for therapeutic intervention in nephrotic kidney disease.

## ACKNOWLEDGMENTS

We thank Dr. Pongpratch Puapatanakul from Chulalongkorn University, Bangkok Hospital, Thailand for his valuable contributions in this study. This work was funded in part by the NSF through the NSF Science and Technology Center for Engineering Mechanobiology (CMMI 1548571), and by the NIH through grants R01DK131177 (to HYS) and R01DK141178 (to JHM and GMG).

## METHODS

### **2.1.** Mouse study and ADR injection

The Adriamycin nephropathy mouse model was used to portray ongoing podocyte injury with similarities to FSGS (Wang, Wang et al. 2000). In brief, male Balb/c mice were injected with ADR (NovaPlus, #NDC0069403112, Stock 2 mg/ml dissolved in 0.9% NaCl pH 3.0, one 15 mg/kg retro-orbital intravenous injection dose) and kidneys were collected at 3, 5-, 7-, 9-, and 14-days post-Adriamycin injection.

### **2.2.** Study Population and Sample Selection

Human kidney 5-μm paraffin sections from patients with nephrotic syndrome (NS) due to minimal change disease (MCD, N=4), primary focal segmental glomerulosclerosis (FSGS, N=5), membranous nephropathy (MN, N=5), and advanced diabetic nephropathy (DN, N=5) were used for this study. Nephrectomy samples (N=7) served as controls.

Formalin-fixed paraffin-embedded (FFPE) de-identified kidney biopsy samples were obtained through the Kidney Translational Research Center (KTRC) at Washington University School of Medicine in St. Louis (approval No. 201102312) and Chulalongkorn University, Bangkok Hospital, Thailand. All samples were archived and spanned the period from 2021 to 2023. Inclusion criteria included the diagnosis of glomerular diseases based on evidence of pathological evidence of the described diseases (i.e., MCD, FSGS, MN, and DN) in the maximally affected glomerulus without immune complex depositions).

### **2.3.** Antibodies and Immunohistochemistry (IHC)

These antibodies were used for immunofluoresence: anti-laminin α5 (Sigma Mouse IgG1 anti-human LAMA5 # AMAB91124) diluted 1:200, anti-nephrin (rabbit monoclonal Clone #2795E anti-Human nephrin from bio-techne R&D Systems) diluted 1:300, anti-synaptopodin IN (American Research Products Guinea Pig anti SYNPO cat# 03-GP94-IN) diluted 1:200, and anti-Myosin IIA (Abnova mouse IgG2b monoclonal anti-Myosin IIA, clone 3C7 # H00004627-M01) diluted 1:200.

FFPE kidney tissue was sectioned into 5-µm slices using a microtome and mounted on glass slides. Deparaffinization and rehydration were carried out through sequential immersion in xylene (2 cycles of 15 minutes), 100% ethanol (2 cycles of 10 minutes), 95% ethanol (3 cycles of 8 minutes), and 70% ethanol (1 cycle of 8 minutes), followed by rinsing twice with distilled water. Epitope retrieval was performed by heating the sections to 95°C in Tris-EDTA (TE) buffer (pH 9.0) for 40 minutes., then placing them on wet ice to cool to 40°C. The sections were subsequently washed with 1X phosphate-buffered saline (PBS) (3 cycles of 3 minutes) and blocked with 2% bovine serum albumin (BSA) in PBS for 30 minutes at room temperature to block nonspecific binding. The slides were then incubated overnight at 4°C with primary antibodies diluted in blocking solution. Afterward, they were washed with PBS, incubated for 45 minutes at room temperature with fluorescence dye-conjugated secondary antibodies, washed again with PBS, and mounted using an anti-fade mounting medium (Invitrogen).

### **2.4.** Expansion microscopy (U-ExM)

U-ExM protocol on kidney sections was performed as previously described (Langner, Puapatanakul et al.). Briefly, deparaffinized and rehydrated FFPE kidney sections were subjected to anchoring solution (Formaldehyde (0.7%)/acrylamide (1%) for 2 hours in room temperature. Next, for gelation, it was replaced by U-ExM monomer solution (19% Sodium acrylate/ 10% Acrylamide/ 0.1% Bis-Acrylamide in 10x PBS) with 0.55% TEMED and APS and incubated overnight in room temperature. Next day, hydrogel dimensions were measured at this point (1^st^ pre-expanded gel measurement step), and hydrogel-embedded tissues were denatured in SDS buffer (200 mM SDS/200 mM NaCl, 50 mM Tris, pH 9) for up to 4 hours in 95°C. Then, hydrogels were washed extensively with ddH2O (at least 5 times, 20 min each). After the final wash, hydrogels were incubated in fresh ddH2O overnight in room temperature. Hydrogel dimensions were measured (2^nd^ post-expansion gel measurement) and ∼4 times expansion was confirmed by comparison with the pre-expansion dimensions.

For staining and imaging, small gel pieces (12mm diameter) were cut out using a biopsy punch, washed with PBS (at least 2 times, 15 min each) to release water and incubated overnight at 4°C with primary antibodies diluted in PBS-BT (PBS containing 3% BSA and 0.1% Triton X-100). After washing the gel punches 3 times with PBST (PBS containing 0.1% Triton X-100), they were incubated with secondary antibodies diluted in PBS-BT for 3 hours in 37°C. Afterwards, they were washed 3 times with PBST, and then twice with ddH2O (30 minutes each time). For imaging, residual water was removed with a Kimwipe and gels were placed in either glass-bottom 24-well plate or 35mm imaging dish. For better attachment and to reduce gel sliding during imaging, dishes were sometimes pre-coated with poly-L-lysine.

### **2.5.** Imaging

Fluorescence super-resolution images were acquired using both Zeiss Airyscan confocal microscope and Nikon N-SPARC. A 60x objective lens was used for all images. Whole-glomerulus views were acquired using 2×2 or 4×4 tiled imaging, enabling precise evaluation of 1– 2 glomeruli per biopsy slide. Five imaging channels, including the Transmitted light (TD) channel, 405, 488 and 647 nm lasers were utilized to assess both fluorescence and tissue morphology. Digital zoom was applied to magnify specific regions of interest. To capture the three-dimensional architecture of selected glomerular structures, Z-stack imaging was performed, and maximum intensity projection (MIP) was then used to generate single composite images that preserve spatial localization and signal intensity across planes.

Nikon N-SPARC platform. This system features super-resolution scanner and fast reading scanner. Equipped with 1x, 20x, 40x, 60x oil, 60x water objectives and lasers allowing to image within four channels: 405, 488, 564 and 647. To visualize the FP structure, 60x oil lense was used while a larger-area image were acquired (2 times zooming) first with 2K resolution, then the *en face* area were identified and crop for the final presentation.

All images were captured under identical settings, deconvoluted, and subjected to minimal processing, including standardized brightness and contrast adjustments, to maintain consistency in fluorescence signal comparisons. Image processing and quantitative analyses were performed using ImageJ (version 1.54f).

### **2.6.** Quantification and statistical analyses of the images to determine SLS stages

For image quantification, three representative, well-defined glomeruli were selected from the glomerular disease groups and three from the control group. Glomeruli were excluded from analysis if they demonstrated artifacts, global sclerosis, or tangential sectioning that interfered with morphometric accuracy. From the remaining glomeruli, three were selected based on preservation and staining quality, prioritizing well-oriented glomerular tufts with robust laminin and podocyte marker expression. This selection strategy helped optimize SLSs measurement accuracy. The Synpo-positive area was segmented by color thresholding on Z-projected images of individual glomeruli to represent total FP areas. Areas of FPE containing SLSs were identified by prominent Synpo expression and loss of nephrin. Co-staining for Synpo and MyoIIA allow visualization of SLSs.

All statistical analyses were performed using GraphPad Prism (version 10.3.1.) and STATA. Summary statistics (mean, median, percentiles, standard deviation, and 95% confidence intervals) were calculated across control and glomerular disease groups (MCD, FSGS, MN and DN). Differences in SLSs were assessed using One-way ANOVA followed by multiple group comparison analysis with Tukey correction (post-ADR injection) or Holm-Šídák’s correction (MCD, FSGS, MN and DN). A p value <0.05 was considered statistically significant and denoted as follows: ****p <0.0001 (versus control or Day3) or ^####^p<0.0001 (versus Day5).

## Notes

### Competing Interest Statement

The authors have declared no competing interest.

