## Supplementary figures and images for "Progressive Actomyosin Remodeling Defining a Structured Podocyte Injury Response in Nephrotic Syndrome"

### Supplementary Figures

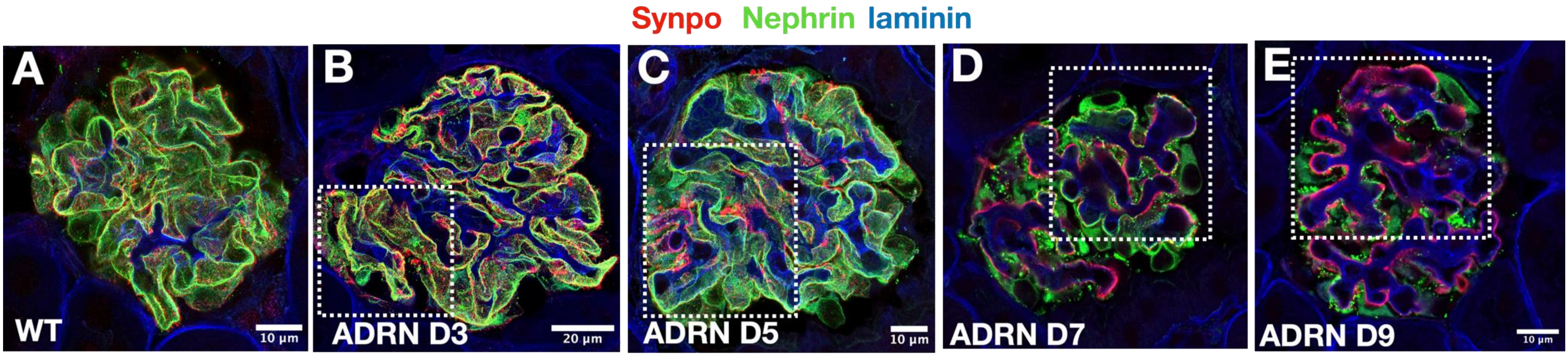

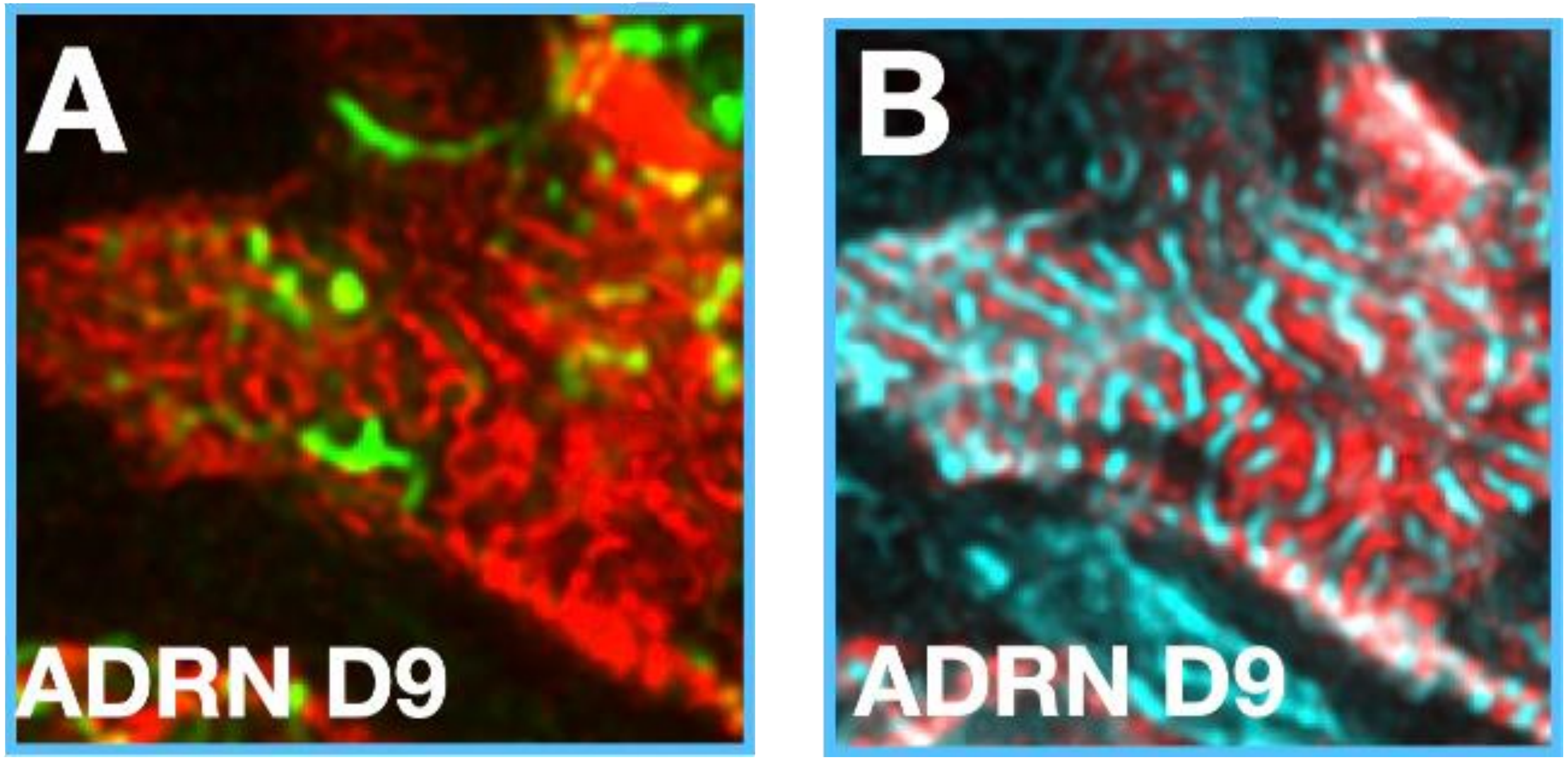


Figure. S2
